# Congruence of location-specific transcriptional programs in intestinal organoids during long-term culture

**DOI:** 10.1101/600940

**Authors:** B. van der Hee, O. Madsen, H. Smidt, J.M. Wells

**Author notes:** Corresponding author: J.M. Wells.

## Abstract

The emergence of intestinal organoids, as a stem cell-based self-renewable model system, has led to many studies on intestinal development and cell-cell signaling. However, potential issues regarding the phenotypic stability and reproducibility of the methodology during culture still needs to be addressed for different organoids. Here we investigated the transcriptomes of intestinal organoids derived from the same pig as well as batch-to-batch variation of organoids derived from different pigs over long-term passage. The set of genes expressed in organoids closely resembled that of the tissue of origin, including location specific functions, for at least 17 passages. Minor differences in gene expression were observed between individual organoid cultures. In contrast, most tissue-specific genes were not expressed in the transformed jejunum cell line IPECJ2, which also showed gene expression consistent with cancer phenotypes. We conclude that intestinal organoids provide a robust and stable model for translational research with clear advantages over transformed cells.

## Introduction

The intestinal epithelium plays an essential role in the digestion and absorption of nutrients while also maintaining homeostasis with symbiotic microbiota [1-3]. The physical containment of microbes to the lumen and homeostasis of tolerance and immunity depends on the functions of different lineages of intestinal epithelial cell [2, 3]. For decades, scientists have exploited the replicative potential of immortalized intestinal cells as enterocyte models to study host-pathogen interactions and intestinal functions *in vitro*. Such monotypic cell models have led to important discoveries but have notable limitations. Immortalized cell lines can undergo significant genotypic alterations within a few passages *in vitro* which are potential threats to data reproducibility [4-6]. Furthermore, transformed cell lines often have altered pathway expression compared to primary cells [4].

Since 2009, when it was shown that intestinal adult leucine-rich repeat-containing G protein-coupled receptor 5 (LGR5+) stem cells [7] could be grown into organotypic cultures and propagated in 3D culture [8, 9], there has been much interest in employing intestinal organoids as advanced models. A distinct advantage of organoids is the development of a crypt-villus axis with a similar spatial organisation of the heterotypic cell lineages found in the tissue of origin. Additionally, methods for generating polarised monolayers of organoids cells [10] have been optimised [11] to improve the versatility of the models, e.g. to study transport, and differential responses to luminal or basolateral stimulants. Another favourable property of organoids generated from adult intestinal stem cells is that they express specific functions associated with their original intestinal location [12]. This means that location-specific functions of different parts of the intestine are intrinsically programmed in adult stem cells.

In the future we can expect intestinal organoids to be increasingly adopted as intestinal models for humans and other mammals. However, there are some unresolved issues that need to be addressed to ensure reliability and reproducibility of results in this emerging field. As organoids contain different cell types there is potential for variability and problems with reproducibility which may compromise their application to phenotype individuals. To address this issue, we assessed the transcriptional stability of intestinal organoids differentiated from the same crypt batch and between organoids from different pigs over long-term passage. Furthermore, we compared expressed genes and pathways in organoids, the original epithelial tissue from which the organoids were derived, and IPEC-J2, a spontaneously transformed porcine cell line derived from the jejunum. The results show that intestinal organoids derived from adult stem cells closely resemble the epithelial tissue of origin in terms of expressed genes and provide a reference for researchers wishing to investigate specific small intestinal functions not present in IPECJ2.

## Methods

### Intestinal organoid generation

Jejunum tissue segments were obtained from control piglets used for another study, following guidelines of the animal ethics committee of Wageningen University. Two 5-week-old piglets were used for generating organoids following procedures previously described [11, 13]. Briefly, a 2 cm section of the mid-jejunum was dissected and placed in ice-cold PBS. After opening the sections longitudinally, jejunum segments were washed three times in ice-cold PBS and villi removed by carefully scraping with a scalpel. Small sections of the mucosa were cut from the muscle layer, divided into small cubes, and transferred into ice-cold PBS containing 30 mM EDTA and incubated with rotation at room temperature for 15 min. The PBS - EDTA was then replaced and incubation continued for 10 min at 37 °C. After washing in ice-cold DMEM supplemented with 5% penicillin/streptomycin (Gibco, Thermo Fisher Scientific), the crypts were dissociated by rigorous vortexing and passed through a 100 µm strainer into cold DMEM containing 5% foetal bovine serum (FBS, v/v). Crypts were pelleted by centrifugation at 300 x g for 5 min, and suspended in Matrigel (Basement Membrane, Growth factor reduced, REF 356231, Corning, Bedford, MA, USA). To improve surface tension, empty 24-well plates were pre-incubated overnight at 37 °C. Matrigel containing crypts was then plated at 7 domes per well (approx. 35 µl per well) and inverted to polymerize at 37 °C. After polymerization, 600 µl F12 cell culture medium (Gibco) was added, supplemented with 100 μg/ml primocin (Invivogen), 10 mM HEPES (HyClone), 1 × B-27 (Gibco), 1.25 mM *N*-acetylcysteine (Sigma), 50 ng/ml human epidermal growth factor (R&D systems), 15 nM gastrin, 10 mM nicotinamide, 10 μM p38 MAPK inhibitor (Sigma), 600 nM TGFβ receptor inhibitor A83-01, and conditioned media for recombinant Noggin (15% v/v), Spondin (15% v/v), and Wnt3A (30% v/v) provided by dr. Kuo and Hubrecht Institute (Utrecht, the Netherlands). Organoids were passaged at a 1:5 ratio every 5 days using ice-cold DMEM by mechanical dissociation, centrifuged at 500 x g for 5 min, and plating in fresh Matrigel matrix droplets as previously described [11].

### Culturing methods and RNA isolation

Directly after crypt isolation, organoid cultures were separated into three batches per animal and grown independently for 17 passages. Jejunum organoids were grown for 3 and 12 weeks (4-17 passages) and extracted using ice-cold PBS. After washing twice in PBS, intact organoids were pelleted at 300 x g for 5 min and suspended in RLT lysis buffer and stored at-80°C prior to RNA isolation. Porcine jejunum epithelial cell line IPEC-J2 (ACC-701) was grown in DMEM F12 medium supplemented with 10% FBS and 5% penicillin/streptomycin (P/S, Gibco) in 75 cm^2^ culture flasks. Data for one IPEC-J2 transcriptome was kindly provided by dr. Richard Crooijmans, via the Functional Annotation of ANimal Genomes (FAANG, BioSamples accession SAMEA4447551)[14]. For the analysis of the remaining two lines, IPEC-J2 at passage 87 and 91 were seeded at 5 x 10^4^ cells/well in 24-well plates and grown to confluence within 48h with reduced P/S (1%). The monolayers were subsequently left to differentiate for 5 days, lysed using RLT-buffer, and stored at −80 °C prior to RNA isolation. Total RNA was extracted using an RNeasy mini-kit (Qiagen) following manufacturer’s instructions including a 15 min on-column DNAse step. Preliminary tRNA concentrations, contamination and degradation were identified using Qubit (Thermo-Fisher) and gel-electrophoresis. The quantity and integrity of RNA was measured using a Bioanalyzer 2100 (Agilent).

### RNA-sequencing procedures and data handling

A minimum of 1 µg total RNA in 50 µl was used for library preparation using the TruSeq RNA sample preparation kit (Illumina) following the manufacturer’s protocol at Novogene. Briefly, total RNA samples were depleted for ribosomal RNA using the RiboZero kit and enriched for mRNA using oligo(dT) beads, fragmented, and synthesized into cDNA using mRNA template and hexamer primers. Custom second strand-synthesis buffer (Illumina), dNTP’s, RNAse H and DNA Polymerase I were added for second strand synthesis initiation. Furthermore, following a series of terminal repair, cDNA library construction was completed with size selection and PCR enrichment. Samples were sequenced using an Illumina Hi-Seq 4000 (Novogene, HongKong) at 9 GB raw data/sample with 150 bp paired-end reads. Raw sequencing reads were checked for quality using FastQC (v0.11.5; [15]) and trimmed using trim-galore for adaptors and quality (v0.4.4 [16]). Only paired-end reads longer than 35 bp were included for further downstream analysis. Sequences were aligned against Ensembl *Sus scrofa* reference genome and annotation 11.1.91 [17] using Tophat (v2.1.1 [18]). Transcriptomes were assembled with 5 bp intron overhang tolerance, merged, normalized, and analysed using the Cufflinks package (v2.2.1 [18]). Differential expression was analysed with 0.01 false discovery rate (FDR) using cuffdiff with bias and weight correction and visualized in R using CummeRbund (v2.7.2 [18]). Mapping analytics can be found in supplementary table S1. Fragments per kilobase million (FPKM) were calculated and log-transformed for downstream analysis. The sequencing data was also processed using CLC Genomics Workbench 11 (Qiagen) using identical reference genome, annotation and settings for identification of insertions/deletions, breakpoints, structural variants, and track generation. Output was filtered for genes <1 FPKM to establish expression.

### Functional analysis and pathway expression

Differentially expressed genes were clustered by k-means into seven clusters using CummeRbund. Due to limited analysis methods for further downstream functional analysis in pig, databases for human genomes were used as a background. Differentially expressed genes were analysed for functional enrichment and ontologies using the TOPPfun suite [19], and gene list enrichment and candidate prioritization were evaluated with a threshold <0.05 for P- and Q-value adjusted for FDR with the Benjamini-Hochberg procedure (B&H). Ingenuity Pathway Analysis (IPA, Qiagen) was used for determining overlapping networks of expressed genes between tissue and organoids, and overall expression of molecular and cellular development of organoids between time-points. Genes for tissue-specific expression were acquired from the human protein atlas for specific mRNA transcription in the small intestine, i.e. duodenum and jejunum, and porcine orthologues were identified to determine tissue and group enriched genes [20].

### Histological analysis

Whole mount imaging was performed as previously described [21]. Paraformaldehyde (PFA; 4%) fixed organoids were stained using FITC-conjugated UEA-1 antibody (FL-1061, Vector Laboratories, USA) and counterstained with Hoechst (0.5 µg/ml, 33342, Thermo Fisher) for 10 min at room temperature. Z-stacks of whole organoids were imaged on a confocal microscope (Zeiss). Immunohistochemical analysis for Mucin-2 was performed following procedures previously described [11, 22]. Organoids were retrieved from Matrigel using ice-cold PBS, pelleted, and fixed overnight using 1% PFA at 4 °C. After pelleting, organoids were suspended in 2% agarose gel, dehydrated and embedded in paraffin blocks. After cutting sections at 5 µm and subsequent drying on glass slides, sections were rehydrated, blocked in 5% normal goat serum, and stained using anti-MUC2 antibody (AB_1950958, GeneTex) overnight. After addition of secondary FITC-antibody (Thermo-Fisher), sections were counterstained using Hoechst and imaged using a DM6 microscope fitted with DFC365 camera at 40x magnification. Generation of chromosomal spreads was achieved following protocols described in [23].

## Results

### Comparative analysis of the transcriptome of organoids, tissue and transformed cells derived from the porcine jejunum

Jejunum tissue was isolated from two euthanized 5-week-old piglets for generating organoids and isolation of RNA from epithelial cells (Figure 1A). Triplicate batches of each organoid were separately cultured for 12 weeks by passaging approximately every 5 days. After 3- and 12-weeks continuous culture (4-17 passages) RNA was isolated from the organoids for RNA sequencing (Figure 1A). Within the first 2 passages after isolation, the organoids acquired a budding phenotype, which might be attributed to whole crypt isolation also containing transit amplifying cells in differentiation stages; as opposed to basal culture after two weeks forming a more cyst-like phenotype. After 2 weeks continuous culture the organoids formed spheroids (Fig 1B) but maintained different cell lineages, as shown with UEA-1 staining for Paneth and Goblet cells (Fig 1C), and a constant chromosome count after 12 weeks (Fig 1D). Similarly, RNA was isolated from the spontaneously transformed porcine jejunum cell line IPEC-J2 at passage number 67, 87 and 91. RNA sequencing data was analysed using a customised analysis pipeline and CLC genomics workbench 11. Multidimensional scaling showed the transcriptomes of the organoids clustered closely together, despite 12-weeks continuous passage and isolation from different pigs (Fig. 2A). It was also evident that the transcriptomes of the tissue and organoid samples were most similar, not separating in the first dimension, whereas IPEC-J2 separated from tissue in both dimensions. A correlation matrix of transcriptomic data from all samples revealed highest similarity for replicate samples of the same origin (Fig 2B).

**Figure 1.**
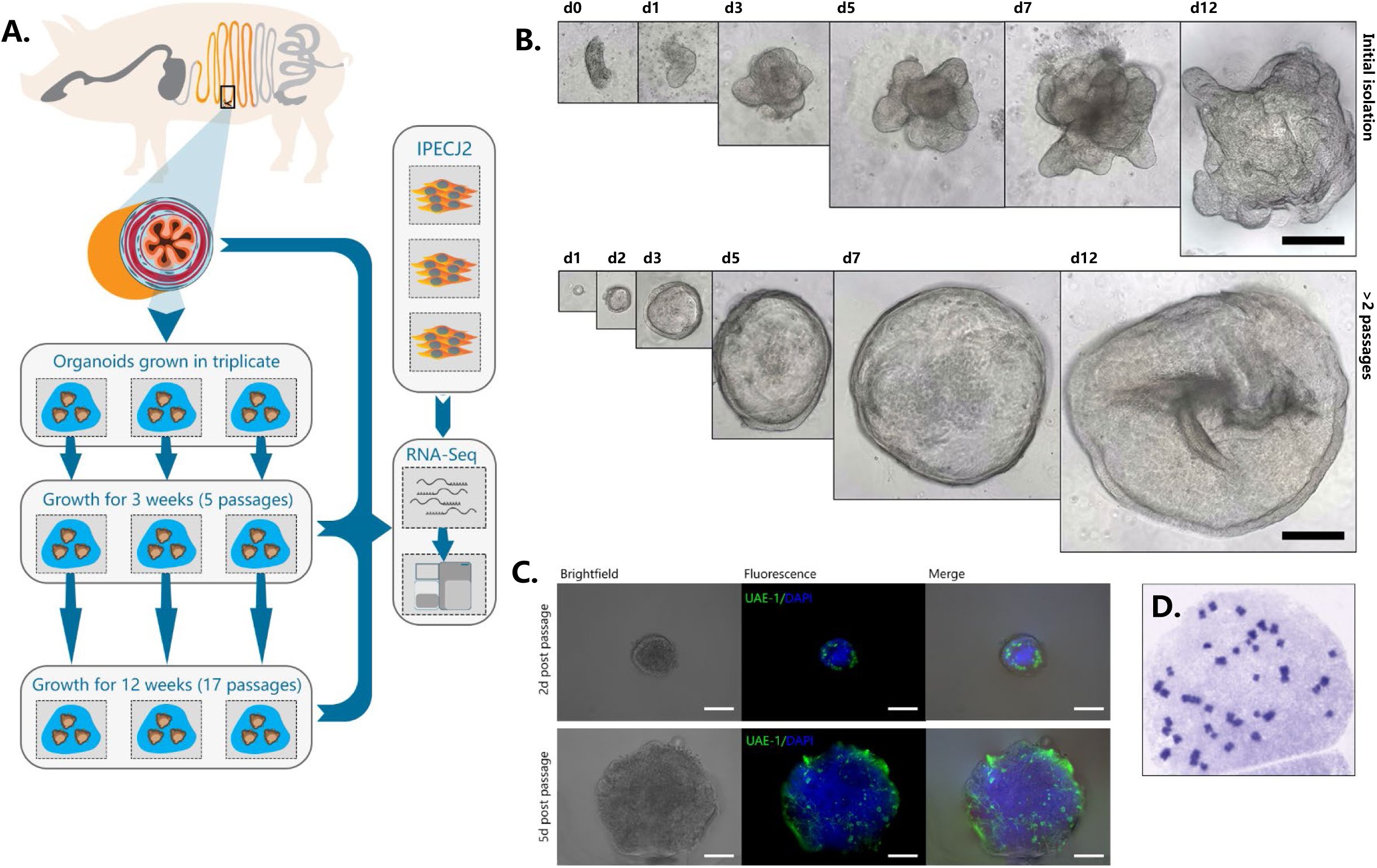
Culture of porcine intestinal organoids and study setup. (A) graphical overview of the study design. Organoids were generated from the jejunum of two individual pigs, directly divided into triplicate organoid cultures per animal, and passaged for 12 weeks. Total RNA of tissue, organoids, and jejunum cell line IPEC-J2 was extracted and sequenced by RNA-seq. (B) Initially after isolation, intestinal crypts form budding organ-like structures *in vitro*. Within two weeks of passaging, organoids form more spheroid-resembling structures for the remainder of the experiment, indicating more long-term reproducibility after a short-term series of passaging. (C) After 12 weeks of growth, spheroids still retain secretory cell lineage differentiation (goblet and Paneth cells) within early post-passaging (green: stained with UEA-1). (D) After 12 weeks of culturing, the organoid chromosomal count was still n = 2×19.

**Figure 2.**
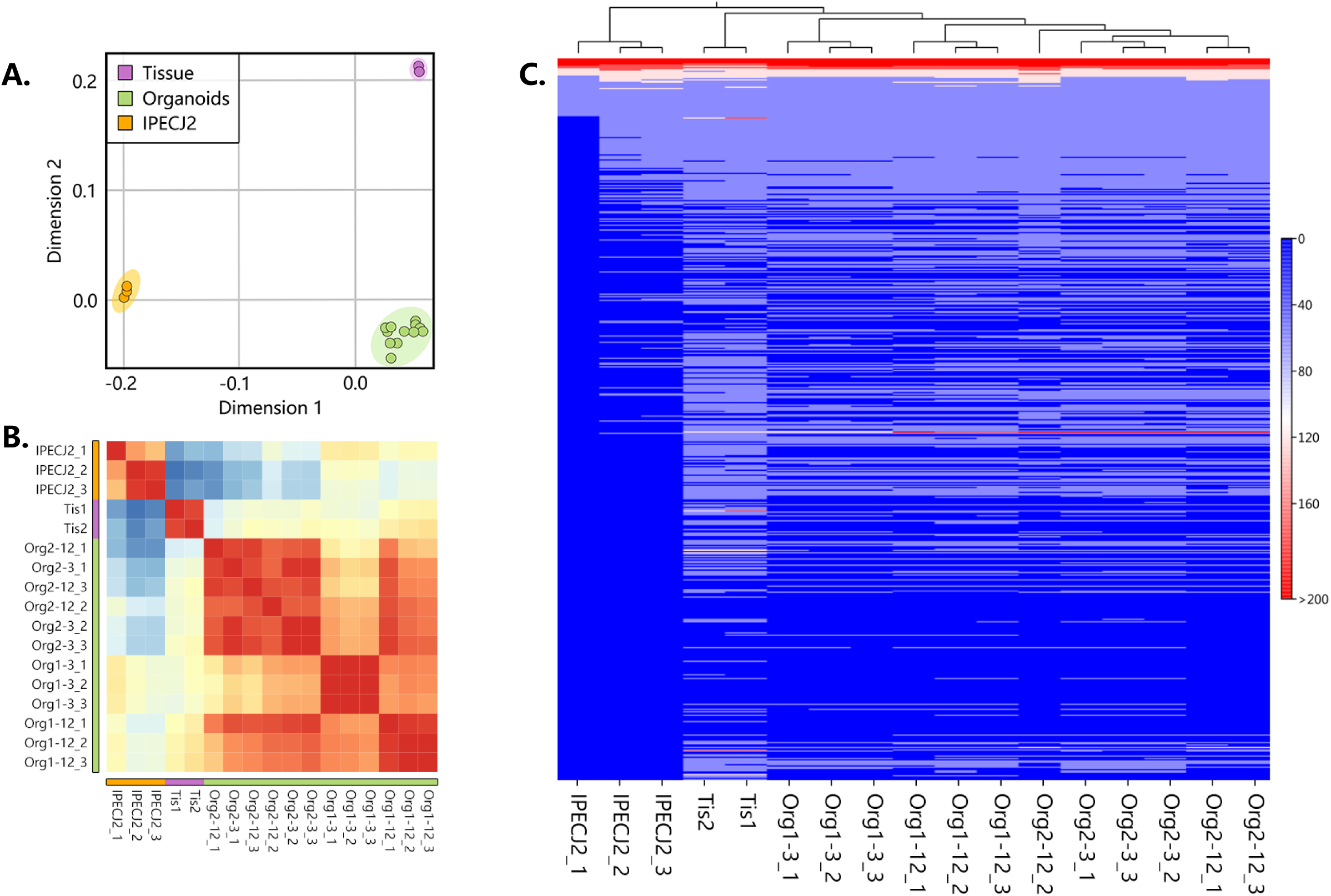
Overview of transcriptome analysis of tissue, organoids, and IPECJ2. (A) Multidimensional scaling of transcriptomic data showed separation of organoids, tissue and IPEC-J2, where organoids and tissue show separation in only one dimension. (B) Correlation matrix of all samples show high correlation among individual organoid batches, and strong correlation between tissue. (C) Heat map and hierarchical clustering of all expressed genes (>1 FPKM) in the dataset.

The hierarchical clustering of gene expression (Fig 2C) further revealed that the transcriptome of organoids more closely resembled that of jejunum epithelial tissue than the IPECJ2 cell line. The two tissue samples clustered under the same branch, where organoids showed consistent homogenous expression derived from different pigs than between 3 and 12-week cultures of replicate samples of the same organoid batch. Furthermore, organoid transcriptomes showed better correlation to tissue (r = 0.77) than IPEC-J2 (r = 0.57), whereas between organoids and IPEC-J2 correlation is higher (r = 0.73) (Suppl Figure 1).

### Organoid transcriptomes closely resemble gene expression signatures associated with their tissue of origin

Of the 24912 annotated genes in the pig genome, 11099 (44.6%) were not expressed in the dataset (<1 FPKM). All samples shared expression of 9117 genes, and a large set of genes was commonly expressed between organoids and tissue exclusively (1762 genes; Fig 3A). Genes associated with different overlapping areas of the Venn diagram were categorized by gene ontology using TOPPfun (Sup table X1). However, the dataset for expressed genes in organoids still contained 1304 genes not annotated denoted with an unknown gene ID, and after conversion to human orthologues using g:profiler [24], only 121 of these unknown ID’s with unknown function acquired a gene annotation (Sup table X2). Nevertheless, after conversion most of the 1304 genes did contain a gene description (80.6%). It is therefore evident that further curation of the porcine ontology database is necessary to generate a more comprehensive reference genome for transcriptomics research.

**Figure 3.**
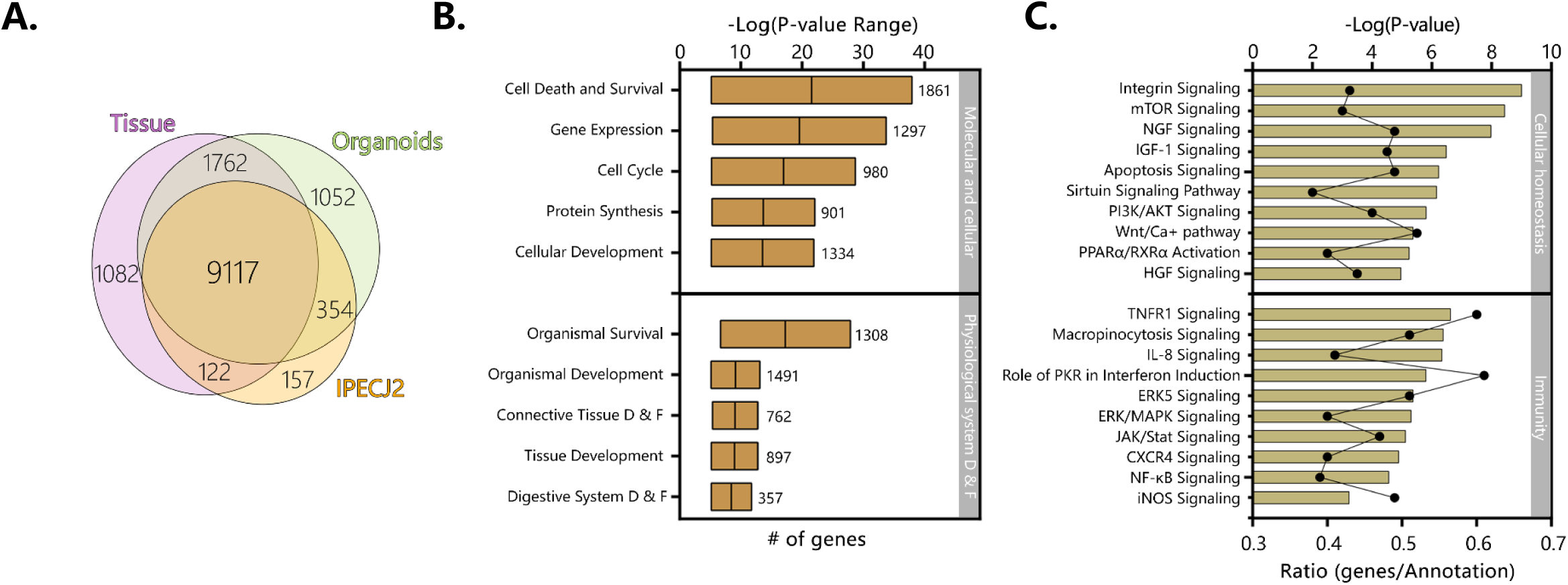
Jejunum organoids are associated to development- or physiology related pathway clusters and show strong overlap with tissue and IPEC-J2. Averages of all expressed genes were compared between sample type and (A) can be viewed in the weighted Venn-diagram. All genes expressed in organoids after 12 weeks of culture were analyzed using Ingenuity pathway analysis. (B) Molecular, cellular, and physiological system development and function shows many genes involved in basic cellular and tissue specific processes. More than >400 pathways were expressed in the organoid RNA-seq dataset. (C) The top 10 cellular homeostasis and immunity related pathways; −logP values indicate statistical probability of pathway expression; ratio, indicates number of expressed genes divided by number of annotated genes in the pathway.

The top clusters of pathways expressed in organoids are involved in basal molecular and cellular function as well as physiological system development, reflecting the interactions with extracellular factors and self-organisation of organoid microanatomy (Fig 3B). Pathways of particular relevance for organoid models such as homeostasis and innate immunity were significantly expressed in organoids. Cellular homeostatic signalling or activation pathways included integrin, mTOR, Sirtuin, PPARα and RXRα, with typically more than 40% of genes in the annotation being expressed (Fig 3C). Immunity-pathways included important cytokine and cytokine receptor signalling pathways (e.g. TNFR1, IL8, JAK/STAT, PKR) and innate immune signalling (NF-kB, ERK/MAPK, iNOS).

Ingenuity pathway analysis of the 1762 expressed genes shared only between organoids and tissues revealed pathways associated with GPCR and Ephrin signalling, which is associated with processes such as cell migration and stem cell differentiation (Fig 4A). Pathways associated with the endocrine functions of cells were also identified including signalling via tryptophan derived melatonin and serotonin. Other pathways specific to organoids and tissue included pathways linked to cytokine, MAP-kinase and other signalling pathways, which are altered in various disease states. Genes encoding complement factors were also specifically expressed in tissue and organoids. Recent studies integrating intestinal transcriptomes deposited in public databases suggest that intestinal expression of complement pathways plays a homeostatic role, being upregulated by inflammatory challenges to control microbial invasion or colonisation (Benis et al., personal communication, and [25]). From this data it is evident that many complement factors are expressed in organoids and tissue (Fig 4B). A key difference between IPEC-J2 and organoid or tissue was the specific expression of hormones associated with Enteroendocrine cells, such as *CCK* which induces secretion of digestive enzymes and *PYY*, a satiety hormone (Fig 4C).

**Figure 4.**
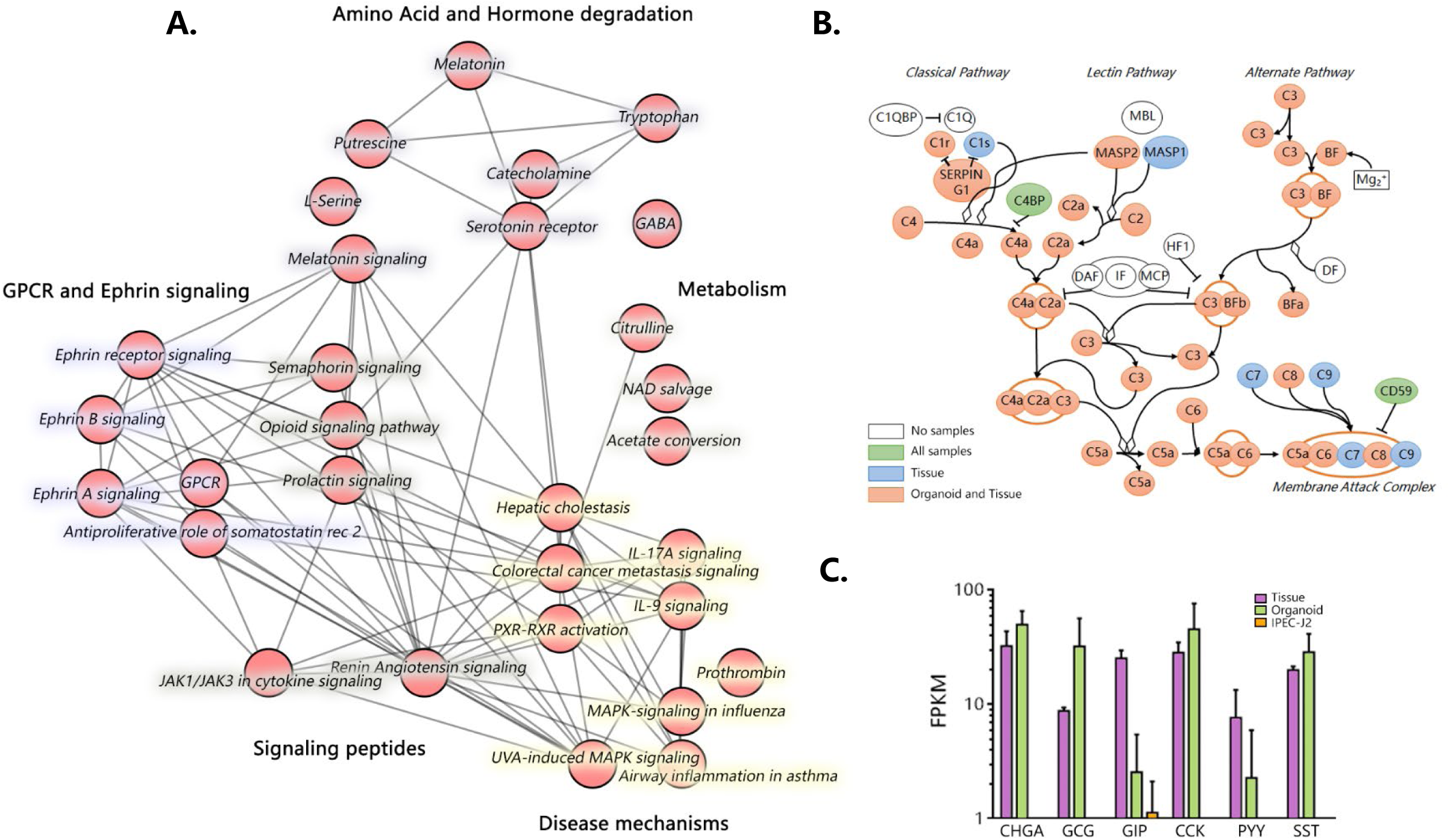
The transcriptome of jejunum organoids exhibits strong similarity to its derived tissue transcriptome, including a distinct group of overlapping genes not expressed in IPECJ2. Testing the RNA-seq dataset for overlapping genes revealed a set of 1762 genes exclusively expressed in tissue and organoids. Top 30 connected canonical pathways of these 1762 genes from ingenuity pathway analysis, which showed subdivision into metabolic, disease, GPCR/Ephrin signaling, and small molecule degradation pathways. (B) Expression of genes involved in the complement pathway are expressed in organoids and tissue (Pink), Tissue only (Blue), organoids tissue and IPEC-J2 (Green), or not found to be expressed (White). (C) Expression patterns of genes involved in Enteroendocrine signaling (CHGA; Chromogranin A, GCG; Glucagon, GIP; Gastric inhibitory polypeptide, CCK; Cholecystokinin, PYY; Peptide YY, SST; Somatostatin, Purple; Tissue, Green; Organoid, Orange; IPEC-J2, data shown as Log(FPKM)).

The genes identified in the human protein atlas to be enriched in the small intestine (i.e. jejunum) [20, 26] were checked for expression in porcine jejunum tissue as well as organoids and IPEC-J2 cells. Most of the porcine orthologues (65% in tissue samples) were indeed expressed in epithelial tissue from the pig jejunum (Fig 5A). Moreover, between 74-86% of the porcine jejunum tissue-specific genes were also expressed in organoids, whereas only 32% were expressed in IPEC-J2 cells (Fig 5A). Genes associated with general digestion (*SI, FABP1*), absorption (solute-carriers; *SLC*-genes), or immunity (*IL22RA1*) were exclusively expressed in the tissue and organoids. Some of the genes associated with secreted peptides for immunity and digestion were not expressed in organoids and IPEC-J2 (i.e. *MEP1A, MEP1B, AQP10, CCL11*). Genes which are specific for the different cell lineages found in the small intestinal epithelium [27], were generally highly expressed in organoids (Fig 5B) but largely absent in IPECJ2. Surprisingly IPEC-J2 lacked expression of some genes commonly associated with mature absorptive enterocytes, even though cultures of this cell line are reported to differentiate into functional epithelium [28].

**Figure 5.**
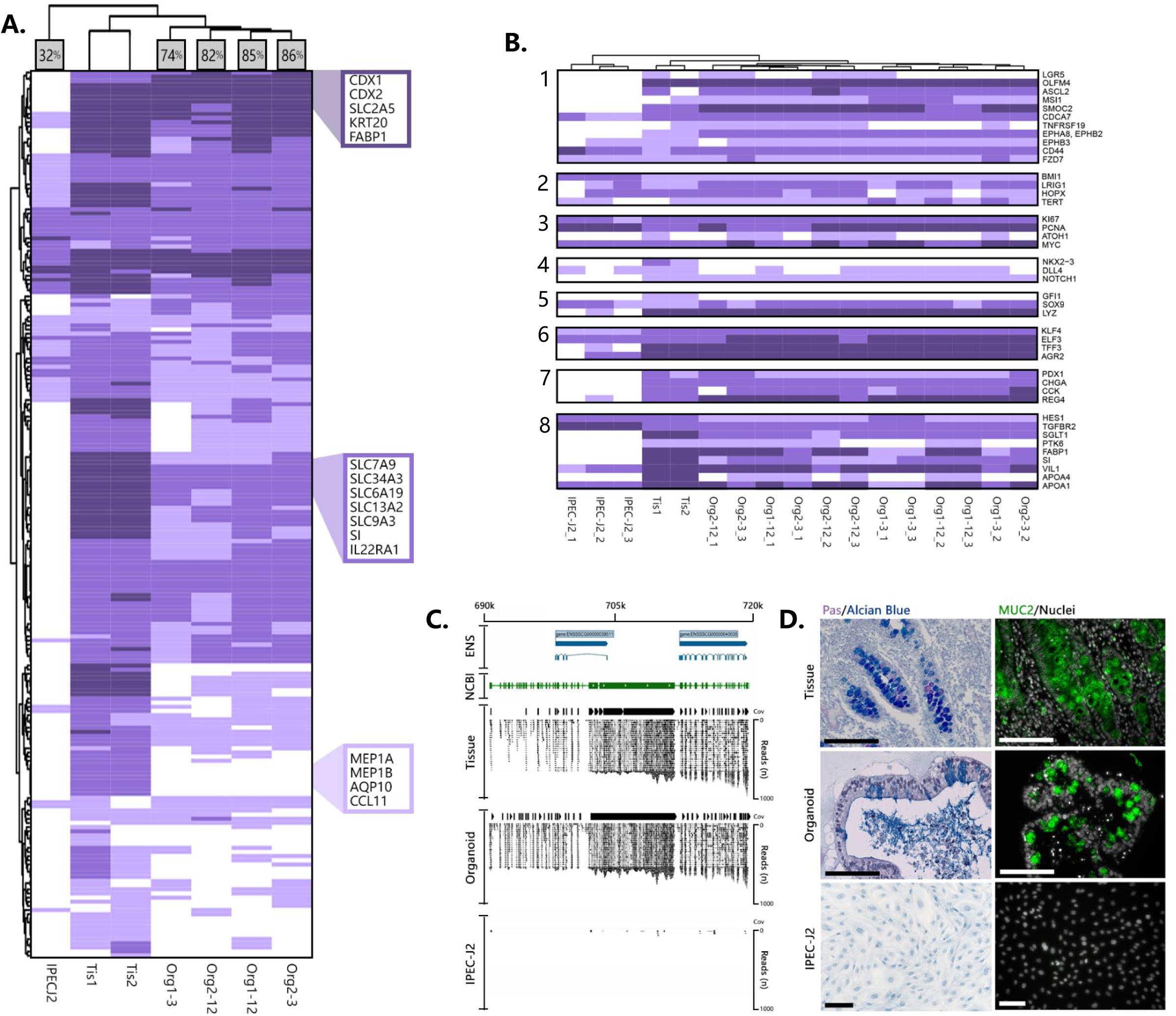
Organoids derived from adult intestinal stem cells are intrinsically programmed to differentiate into different epithelial cell linages and express location-specific transcriptional programs. (A) the majority of jejunum-specific genes identified in the human protein atlas are also expressed in porcine tissue and porcine organoids (>74%), whereas IPECJ2 only expressed 32% of tissue specific genes. (B) Cell type-specific transcripts for (1) Crypt Base Columnar *(CBC)* and Stem cells, (2) Label-retaining *(LRC)* +4 cells, (3) Proliferation, (4) Niche factors, (5) Paneth cells, (6) Goblet cells, (7) Enteroendocrine cells, and (8) Absorptive cells or enterocytes. White: no expression, tones of purple; light (1-10 FPKM), medium (10-100 FPKM), and dark (>100 FPKM) expression. (C) Overlaying the NCBI gene tracks of MUC2 NC_010444.4 on chromosome 2 at location 689363-719542bp (yellow area), shows identical overlap with the mapped reads and coverage (Cov) from organoid and tissue samples, but not in IPECJ2 (ENS; Ensembl reference genome). (D) To confirm MUC2 protein translation and subsequent mucus formation, Carnoy fixed tissue, organoid, and IPECJ2 samples were stained with PAS/Alcian blue (left) and porcine anti-MUC2 (right; black and white size bars indicate 100 µm).

Initially our dataset suggested lack of mucin 2 (*MUC2*) gene expression using Ensembl gene annotation. However, we identified a high number of RNA-seq reads from organoids and tissue mapping to the chromosomal locus associated with *MUC2* in the NCBI database (XM_021082584, Chromosome 2, bp 689,363-719,542) (Fig 5C). We verified the expression of *MUC2* in porcine organoids and tissue using histology confirming mucin production and secretion as described previously [11, 13] (Fig 5D).

### Cluster analysis of differentially expressed genes (DEGs)

Genes which were differentially expressed in tissue, organoids or IPEC-J2 (P<0.05), were categorized by K means clustering and represented as biological processes and pathways (Fig 6A-C). The genes expressed only in tissue (n = 141) are mostly involved in immune pathways e.g. T cell and leukocyte functions, suggesting they might be due to presence of lamina propria immune cells in the tissue sample (Fig 6A). Genes related to extracellular matrix or muscle contraction pathways were also differentially expressed in tissue compared to organoids and IPEC-J2.

**Figure 6.**
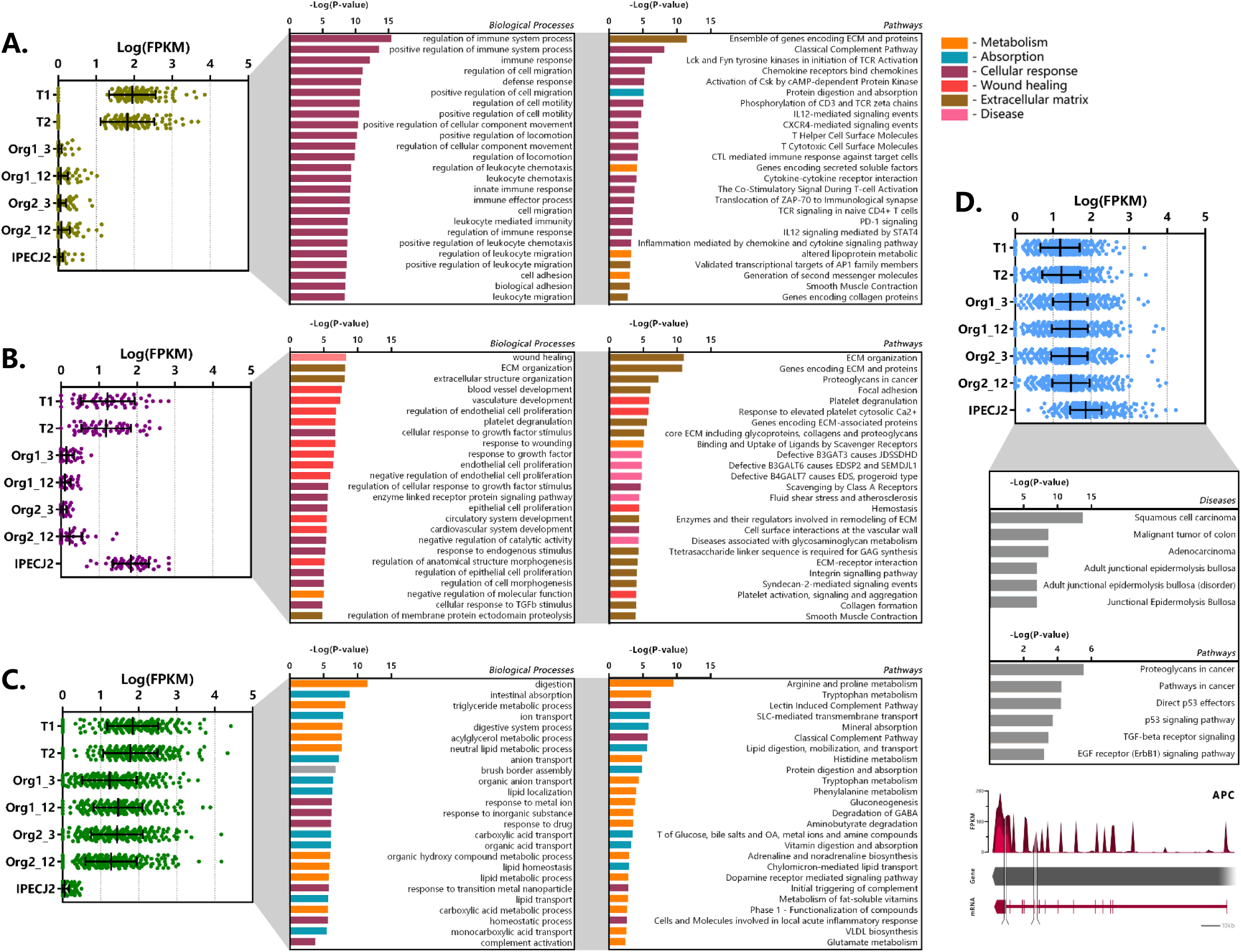
Cluster analysis of differentially expressed genes (DEGs). DEGs were clustered according to their expression pattern using k-means clustering, and subsequently analyzed for functional enrichment using TOPPfun. (A) Cluster of genes (n = 141) showing increased expression in tissue only and low expression in organoids and IPEC-J2. (B) Cluster of genes (n = 52) showing high expression in tissue and IPEC-J2, with low expression in organoids. (C) Cluster of genes (n = 299) with increased expression in tissue and organoids, with low expression in IPECJ-2 (data represented in Log(FPKM) or −Log(P-value), P and q <0.05). (D) Cluster of genes (n = 841) expressed more highly in IPEC-J2 than tissue and organoids which were associated with diseases and pathways involved in tumor formation. A common mutation in colon cancer is inactivation of adenomatous polyposis coli (APC) gene where IPEC-J2 shows an insertion (217 bp, tandem duplication) in the protein coding region and splice site deletion (331 bp, cross mapped breakpoints; bottom).

The cluster of genes expressed at higher levels in tissue and IPEC-J2 (n = 52) compared to organoids comprise of processes related to cell morphology, proliferation, movement, remodelling of the extracellular matrix (ECM) as well as integrin and ECM signalling pathways (Fig 5B). Altered expression of some of these pathways has been reported in cancer [29, 30]. The low or absent expression of the ECM genes in organoids may be due to provision of Matrigel acting as a basement membrane substrate. Furthermore, there may be components present in the intestinal ECM capable of inducing cell integrin signalling which are not present in Matrigel. Clustering genes transcribed in significantly higher amounts in IPEC-J2 show gene-list enrichment in disease processes and pathways associated with cancer (Fig 6D). Furthermore, one of our IPEC-J2 cultures showed an insertion in the protein coding region and a large deletion in the splicing site of the adenomatous polyposis coli (*APC*) gene, which is a tumor suppressor often inactivated in colon cancer [31].

The cluster of genes with higher expression in organoids and tissue (n = 299) compared to IPEC-J2 involve nutrient transport and metabolic processes such as lipid digestion and transport, protein digestion and amino acid metabolism (Fig. 6C). Examples include metabolism of tryptophan, arginine, proline, histidine, phenylalanine and transport of glucose, bile salts, fatty acids, lipids and vitamins.

### Congruence of the transcriptome in individual organoids during passage

Comparison of transcriptomes of organoid cultures over time and between batches indicated high correlation in expression values (r = 0.906-0.910, Fig 7A). After long-term passage there were 199 genes in organoid 1 and 172 genes in organoid 2 which were significantly increased in expression (Fig 7B; n=3 per group). All differentially expressed genes were distributed across 9 common ontologies (Fig 7B) and consisted of only 1.93 to 3.67 % of the annotated genes in each biological process. This included processes such as small molecule biosynthesis, organic acid metabolic processes and response to nutrient levels. Variation in expression of genes in organoid cultures over time may therefore be due to differences in nutrient abundance in culture medium, and replicative activity, rather than permanent loss or gain of functions. Long-term passage also resulted in differential down regulation of 78 and 137 genes in organoids 1 and 2, respectively (n=3 per group, Fig 7C). The only common ontology found for these genes was small molecule biosynthetic process, which was also included in genes upregulated after long term passage (Fig 6B). This suggests that culture dependent conditions result in variation in a small percentage of genes in this ontology group, perhaps due to variation in number or activity of several cell types in low abundance. The heatmap in Fig 7D shows expression or fold change in genes that were commonly differentially expressed in organoids 1 and 2 after long-term passage. Although being significantly differentially regulated, a group of genes with significantly reduced fold change in expression after 12 weeks passage was seen to be overall reduced in transcript abundance (FPKM) and appeared to be involved in unrelated processes when tested for gene ontology. Differences between organoid 1 and 2 are likely to reflect individual variation and the most striking differences were for apolipoprotein A1 (*APOA1*), *ISG12*(A) and *ISG15*.

**Figure 7.**
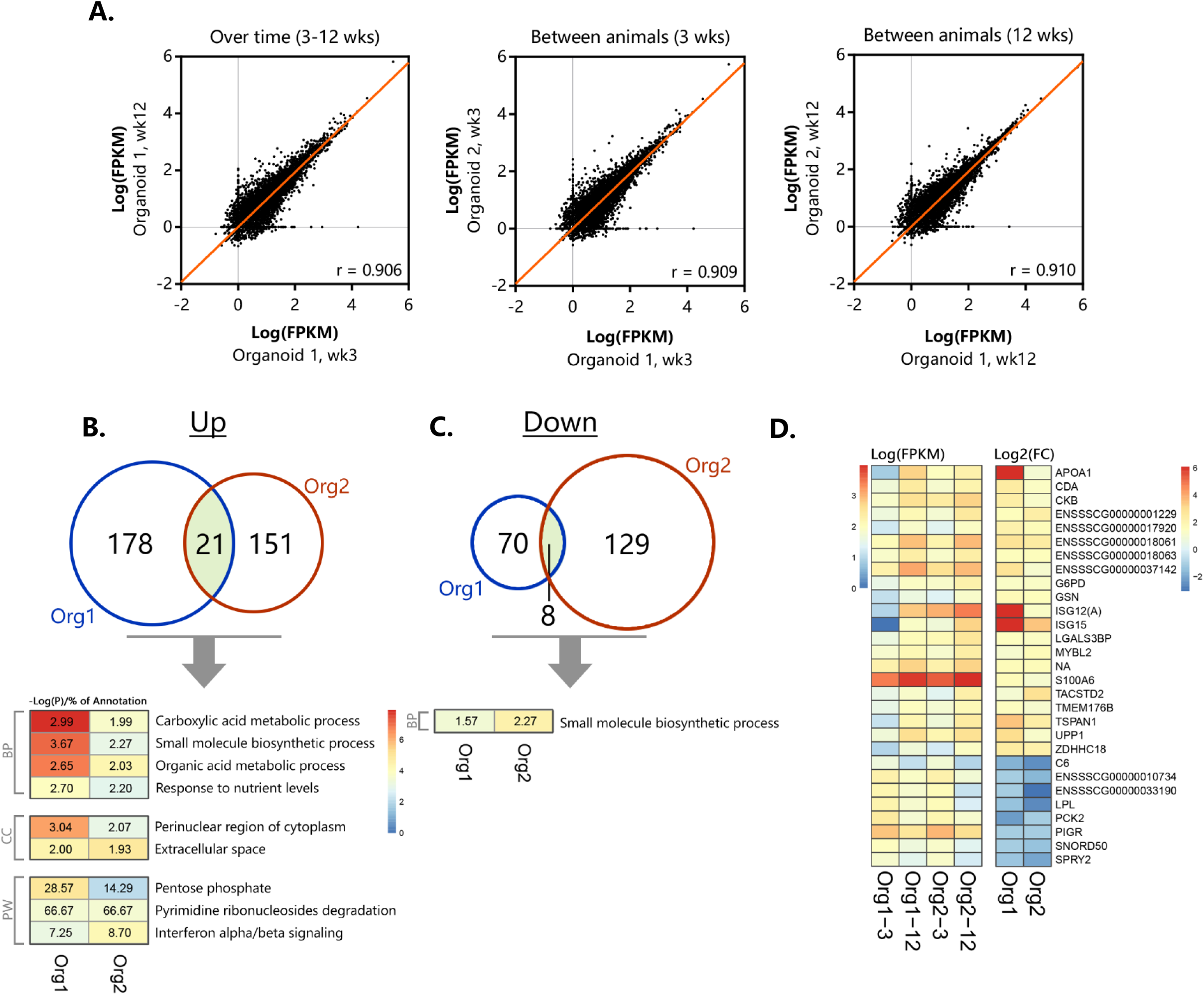
Differential gene- and pathway expression in organoids shows stable transcription over time. The two separate organoid cultures (n = 3 per group) were tested for expression differences between 3 and 12 weeks of culturing, (A) showing representative correlations over time within an organoid line and between different animals (n = 13813 genes with at least 1 sample > 1 FPKM). Significant (p <0.05) up- (B) and down-regulation (C) of genes and their corresponding overlapping pathways when tested individually using TOPPfun (overlapping area shows similar genes between organoid types, BP; Biological processes, CC; Cellular component, PW; Pathway). (D) Overlapping up- and down-regulated genes between 3 and 12 weeks of culturing in actual expression values (Log(FPKM)) and their corresponding fold change (Log2(FC)).

## Discussion

Transformed and cancer cell lines commonly display aneuploidy and mutations affecting cellular physiology, which may further change during long-term passage. Intestinal organoids are considered an attractive alternative to transformed cell lines, but there are no studies evaluating the variability of the transcriptome in different batches of organoids generated from the same crypts and the effect of long-term culture on genomic stability and the transcriptome.

In this study, we compared gene expression in the porcine transformed jejunum cell line IPECJ2, organoids derived from adult stems cells of the porcine jejunum and the intestinal tissue from which they were derived. The genes highly expressed in tissue, but not organoids or IPEC-J2, were mostly involved in immune cell, extracellular matrix or muscle contraction pathways. This is most likely due to the presence of elements of the (sub)mucosa, including lamina propria immune cells in the tissue sample. Porcine intestinal organoids possessed the different epithelial cell lineages found in tissue for at least 17 passages without apparent changes to the karyotype. An interesting observation in organoids was the high expression of *LYZ* which is often associated with Paneth cells. Until recently Paneth cells were considered to be absent in the porcine small intestine [32, 33]. Paneth cells facilitate regeneration of the intestinal mucosa after metaplasia and produce high amounts of *LYZ* in presence of intestinal disease or damage [34, 35].

The set of genes expressed in organoids closely resembled that of the tissue of origin, a characteristic reported for other types of organoids [36, 37]. Moreover, organoids expressed the majority of gene orthologues (>74%) associated with jejunum specific functions in humans for at least 17 passages, showing that the adult stem cells retain their location-specific functions over long-term culture [12, 38].

Overall the batch to batch variation in organoids from the same animal was low, which may have been aided by concurrent passage and consequently differentiation state. The functions of genes which significantly altered expression between organoids from different animals or different cultures suggests they arise from differences in abundance of nutrients in culture medium or replicative activity, rather than permanent loss or gain of functions.

The main described benefits of the jejunum cell line IPECJ2 are its non-cancerous origin and stability for more than 98 passages [28, 39]. However, we observed multiple indications of increased expression of genes associated with cancer or tumors in IPECJ2 (passage 67-91). For example, these included *ANXA1* and *CALD1* [40], as well as insertions and deletions in the tumor suppressor *APC* [31]. We conclude that porcine jejunum organoids more closely resemble jejunum tissue than IPECJ2 and provide a robust model for gene expression studies for at least 12 weeks of culture. As such they provide an advanced model for mechanistic studies on host-microbe interactions and intestinal physiology [41]. Organoids are also likely to avoid changes in glycosylation patterns seen in cancer or transformed cell models. The RNA-seq data provides a valuable resource for researchers to assess the suitability of intestinal organoids for studying specific pathways or biological processes.

Pig organoids can be rapidly generated from left over slaughter material without requirement for ethical approval. Thus, they have potential to be used to identify new phenotypes and investigate the role of genetic polymorphisms in susceptibility to enteric infections or other production traits. Furthermore, porcine intestinal physiology is considered to closely resemble that of humans increasing the potential for translation of results to humans. Moreover, recent progress in developing robust methods for generating 2D monolayers from 3D organoid cultures facilitates apical exposure to test substances and also opens up possibilities for studying epithelial transport [11, 42].

## Acknowledgements

We would like to thank Anja Taverne-Thiele, Nico Taverne, and Linda Loonen for their practical assistance and helpful discussions. Furthermore, we appreciate the provision of one IPEC-J2 RNA-seq dataset (BioSamples database: SAMEA4447551) by dr. Richard Crooijmans, Animal Breeding and Genetics, Wageningen University, The Netherlands, under the FAANG data sharing agreement (http://data.faang.org/home). We thank dr. Sabine Middendorp, Wilhelmina Children’s Hospital Utrecht, The Netherlands, for initial guidance setting up organoid cultures. We would like to thank dr. Kuo, Stanford University, United States of America, for providing the R-Spondin 1 transfected cell line for conditioned medium. Cell lines for Noggin and WNT3A conditioned medium were provided by the Hubrecht Institute, Utrecht, The Netherlands. We thank dr. Arie Kies for critically reading our manuscript and helpful suggestions. This research is financed by the Applied and Engineering Sciences Division of the Netherlands Organization for Scientific Research NWO (project number 14935), the Ministry of Economic Affairs and DSM Nutritional Products. The financial sponsors played no decisive role in collection, analysis, and interpretation of the data, or the decision to submit for publication.

## Supplemental information

**Supplementary Figure 1.**
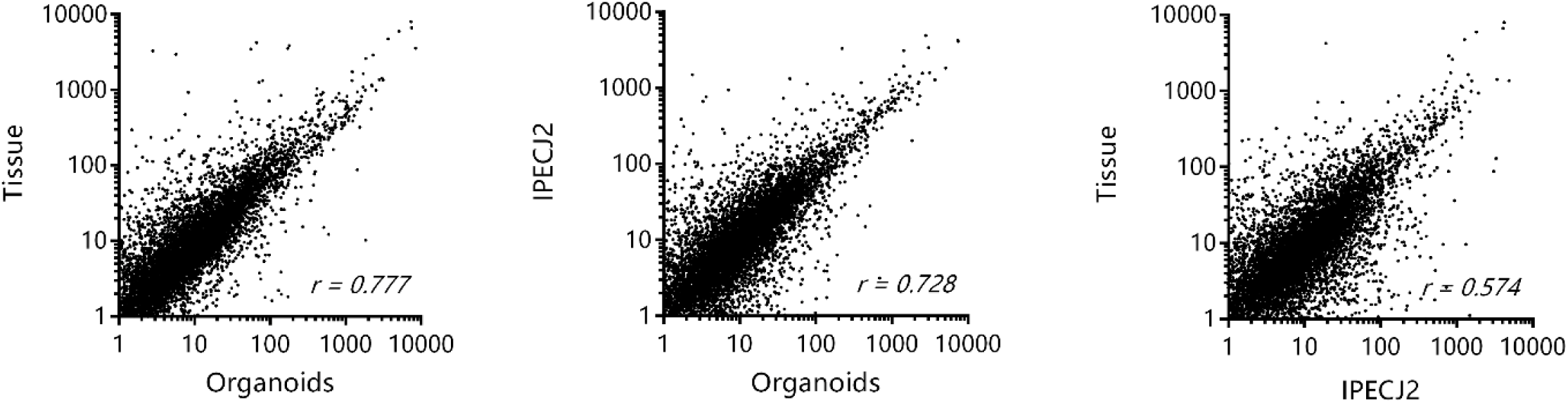
Correlation plot of average FPKM per sample type. (A) Organoids and tissue show strong correlation (r = 0.777), as well as between (B) Organoids and IPEC-J2 (r = 0.728). The least amount of correlation was observed between (C) Tissue and IPEC-J2 (r = 0.574)

**Supplementary table 1.**
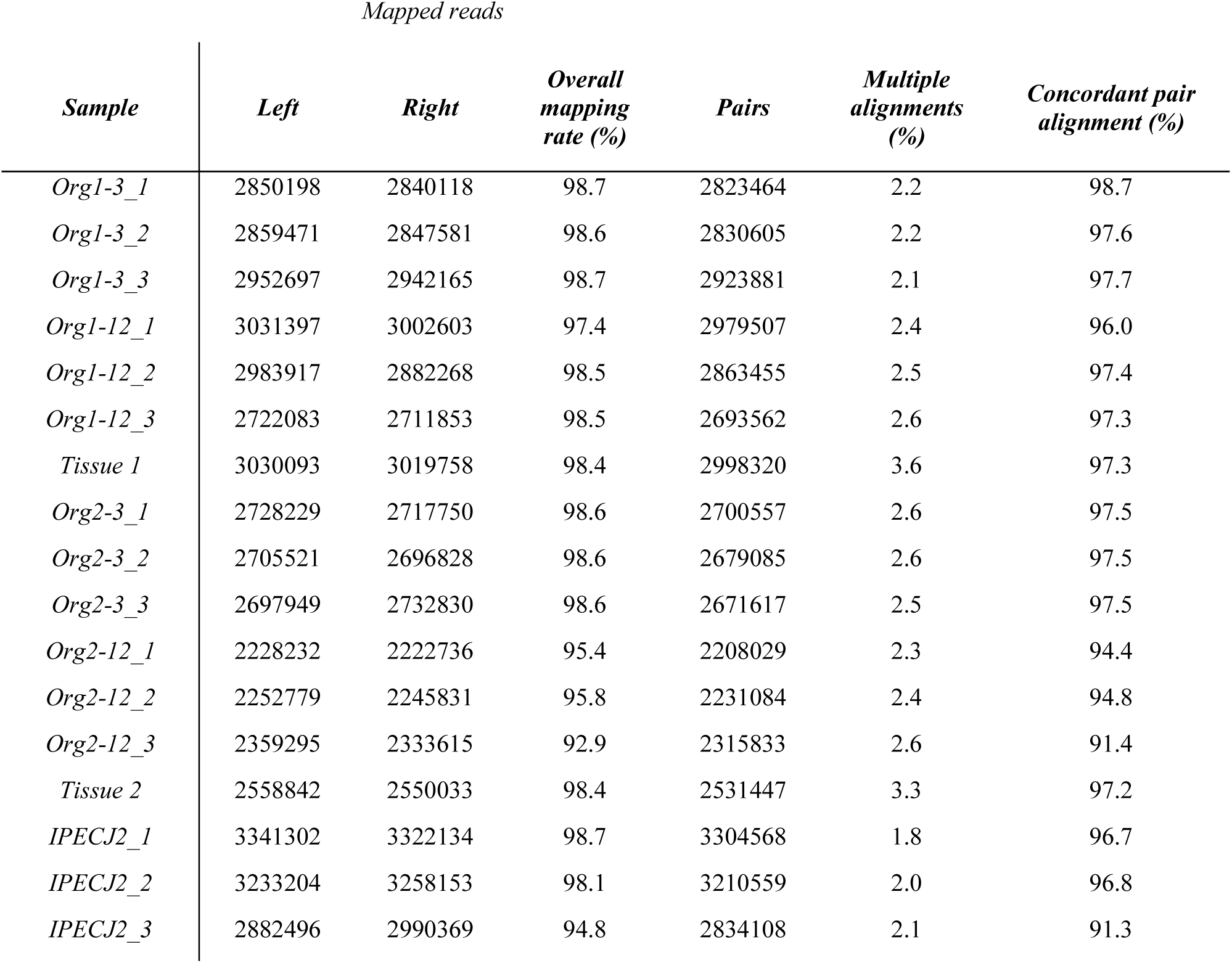
Summary of mapping RNA-seq data to reference genome Sus Scrofa 11.1.

| Sample | Mapped reads |  |  |  |  |  |
| --- | --- | --- | --- | --- | --- | --- |
|  | Left | Right | Overall mapping rate (%) | Pairs | Multiple alignments (%) | Concordant pair alignment (%) |
| *Org1-3_1* | 2850198 | 2840118 | 98.7 | 2823464 | 2.2 | 98.7 |
| *Org1-3_2* | 2859471 | 2847581 | 98.6 | 2830605 | 2.2 | 97.6 |
| *Org1-3_3* | 2952697 | 2942165 | 98.7 | 2923881 | 2.1 | 97.7 |
| *Org1-12_1* | 3031397 | 3002603 | 97.4 | 2979507 | 2.4 | 96.0 |
| *Org1-12_2* | 2983917 | 2882268 | 98.5 | 2863455 | 2.5 | 97.4 |
| *Org1-12_3* | 2722083 | 2711853 | 98.5 | 2693562 | 2.6 | 97.3 |
| *Tissue 1* | 3030093 | 3019758 | 98.4 | 2998320 | 3.6 | 97.3 |
| *Org2-3_1* | 2728229 | 2717750 | 98.6 | 2700557 | 2.6 | 97.5 |
| *Org2-3_2* | 2705521 | 2696828 | 98.6 | 2679085 | 2.6 | 97.5 |
| *Org2-3_3* | 2697949 | 2732830 | 98.6 | 2671617 | 2.5 | 97.5 |
| *Org2-12_1* | 2228232 | 2222736 | 95.4 | 2208029 | 2.3 | 94.4 |
| *Org2-12_2* | 2252779 | 2245831 | 95.8 | 2231084 | 2.4 | 94.8 |
| *Org2-12_3* | 2359295 | 2333615 | 92.9 | 2315833 | 2.6 | 91.4 |
| *Tissue 2* | 2558842 | 2550033 | 98.4 | 2531447 | 3.3 | 97.2 |
| *IPECJ2_1* | 3341302 | 3322134 | 98.7 | 3304568 | 1.8 | 96.7 |
| *IPECJ2_2* | 3233204 | 3258153 | 98.1 | 3210559 | 2.0 | 96.8 |
| *IPECJ2_3* | 2882496 | 2990369 | 94.8 | 2834108 | 2.1 | 91.3 |

